# Guiding dose selection of monoclonal antibodies using a new parameter (AFTIR) for characterizing ligand binding systems

**DOI:** 10.1101/432500

**Authors:** Sameed Ahmed, Miandra Ellis, Hongshan Li, Luca Pallucchini, Andrew M. Stein

**Affiliations:** Department of Applied Mathematics, University of Waterloo; School of Mathematical and Statistical Sciences, Arizona State University; Department of Mathematics, Purdue University; Department of Mathematics, Temple University; Novartis Institute for BioMedical Research, 45 Sidney St, Cambridge MA, 02140 USA

**Keywords:** monoclonal antibody, target mediated drug disposition, pharmacokinetics, pharmacodynamics, pharmacometrics

## Abstract

Guiding the dose selection for monoclonal antibody oncology drugs is often done using methods for predicting the receptor occupancy of the drug in the tumor. In this manuscript, previous work on characterizing target inhibition at steady state using the AFIR metric [1] is extended to include a “target-tissue” compartment and the shedding of membrane-bound targets. A new potency metric AFTIR (Averarge Free Tissue target to Initial target ratio at steady state) is derived, and it depends on only four key quantities: the equilibrium binding constant, the fold-change in target expression at steady state after binding to drug, the biodistribution of target from circulation to target tissue, and the average drug concentration in circulation. The AFTIR metric is useful for guiding dose selection, for efficiently performing sensitivity analyses, and for building intuition for more complex target mediated drug disposition models. In particular, reducing the complex, physiological model to four key parameters needed to predict target inhibition helps to highlight specific parameters that are the most important to estimate in future experiments to guide drug development.

## Introduction

During biologic drug development, prediction of target engagement at the site of action plays a critical role in dose regimen selection [2]. Because target engagement measurements at the site of action are often impossible to obtain, model-based predictions of target engagement at the site of action are often used to help justify the dose regimen selection. The methods used to predict target engagement vary significantly in their assumptions and in their level of complexity. For example, consider the model-based dose selection of pembrolizumab and atezolizumab.

For the PD-1 inhibitor pembrolizumab, a physiologically based model for antibody distribution and target engagement was developed to predict the dose needed to achieve target engagement and tumor suppression [3]. This model involved many assumptions about a large number of parameters and how these parameters scale from mice to humans. For the PD-L1 inhibitor atezolizumab, a much simpler approach was taken to guide the dosing regimen [4]. The tumor biodistribution coefficient (*B*) and in vivo binding affinity (*K*_d_) were chosen based on preclinical data. The steady state trough concentration (*C*_trough_) was estimated from clinical observations. The receptor occupancy (*RO*) formula in Equation 1 was then used to identify the dosing regimen needed to achieve 95% target occupancy. This approach made fewer assumptions about model parameters than in [3]. However, in choosing this simple model, many implicit assumptions were made about the system. For example, this simple model assumes that the rate of PD-L1 internalization does not change when it is bound to atezolizumab.

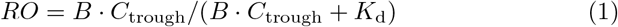

The complex, mechanistic model used for pembrolizumab, captures more details of the underlying physiological processes. However, this model was more difficult to analyze due to it’s complexity; it also had a large number of unknown parameters which were difficult to estimate accurately. The simple model used for atezolizumab was easier to analyize and had fewer unknown parameters to estimate. However, this model required certain implicit assumptions which do not necessarily hold.

In this paper, a mathematical analysis of a physiologically-based model for drug distribution and target turnover is performed to derive a simple potency factor for characterizing target engagement. This theoretical calculation is validated using simulations of the model. All assumptions made in deriving this formula are explicitly stated. This paper extends previous work that focused on target engagement in circulation, as characterized by the Average Free target to Initial target Ratio (AFIR) at steady state in circulation [1].

## Theory

In this section, an expression for the Average Free Tissue target to Initial target Ratio in tissue (AFTIR) is derived for the model in Figure 1.

**Fig. 1.**
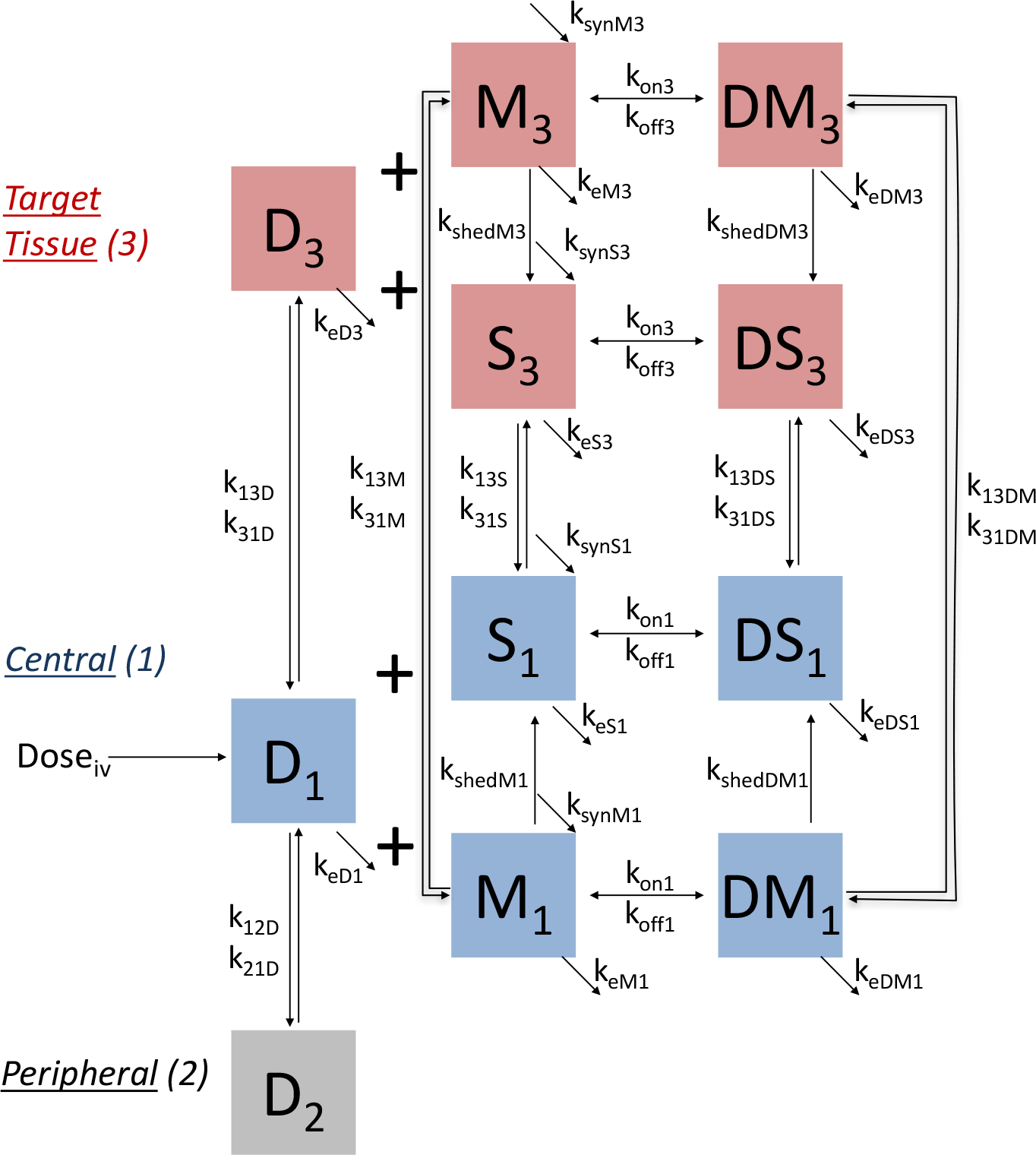
The extended target mediated drug disposition model. Vertical arrows represent distribution, horizontal arrows represent dosing and binding, and diagonal arrows represent synthesis and elimination.

### Model Description

The model in Figure 1 is based on the standard target mediated drug dispositon (TMDD) model [5], where a drug (*D*) binds its target. The model is extended to include the following processes:

– Shedding of the membrane-bound target (*M*) to form soluble target (*S*).
– Binding of the drug to both the membrane-bound target and the soluble target to form complexes *DM* and *DS*, respectively.
– Distribution of drug, target, and complexes from central (1) to tissue (3) compartment via either passive processes (soluble target) or active processes such as cell-trafficking (membrane-bound target).

As with the standard TMDD model, the peripheral compartment (2) contains the drug only; it is included so that the two-compartment pharmacokinetics of the drug can be described. The ordinary differential equations for describing this system are given by Equations (2) – (12).

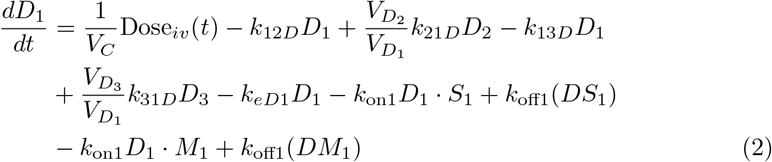

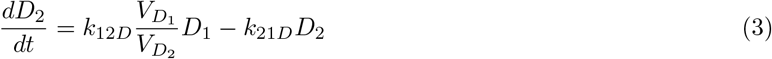

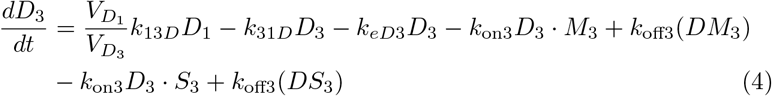

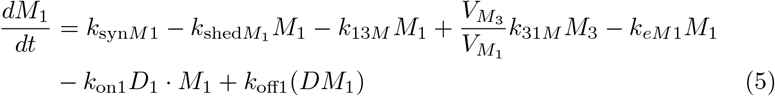

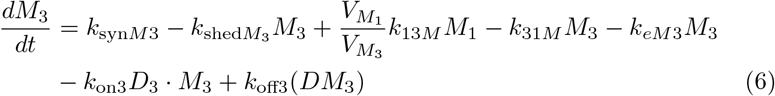

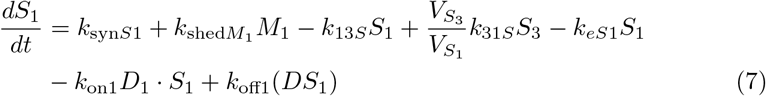

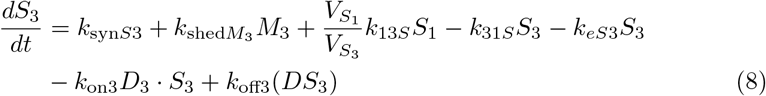

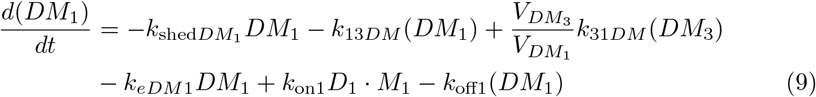

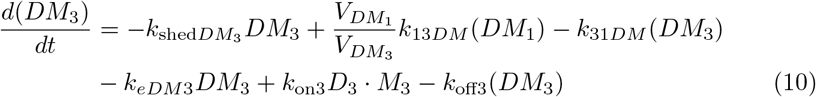

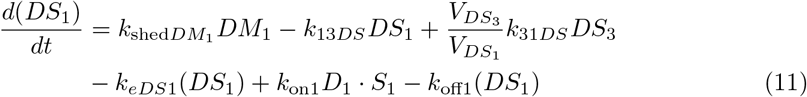

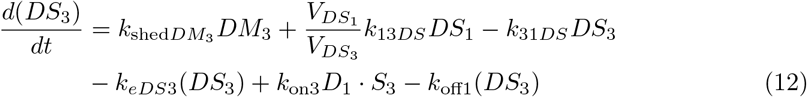

The initial conditions for all variables are zero, except for the free target concentrations *M*_1_, *M*_3_, *S*_1_, and *S*_3_. The equations used to determine these four initial concentrations are provided in Appendix A.1.1.

### Checking for tumor homogeneity using the Thiele Modulus

In setting up the compartmental model in Figure 1, an implicit assumption is made that all compartments can be treated as homogeneous, well-mixed tissues. This assumption is reasonable for the circulating compartment, and it is not needed for the peripheral compartment, which is an empirical model feature that is used to get the standard 2-compartment PK. But when the target is membrane-bound, then for the tissue compartment, this assumption will not be correct for low doses where the drug may be internalized and eliminated before it penetrates through the tissue. For membrane-bound targets, the assumption of a homogeneous, well-mixed tissue compartment can be evaluated by checking whether the Thiele modulus given below is less than one. [6, 7]

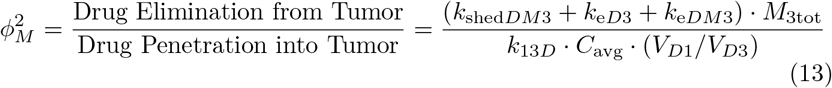

The Thiele Modulus is a non-dimensional parameter which is the ratio of receptor elimination (due to shedding and internalization) to the drug penetration into the tissue. Here, *M*_3tot_ is the total concentration of receptors in the tissue at steady state, which is often assumed to be equal to *M*_30_, but could also be set to *M*_30_ *T*_fold_, where *T*_fold_ is the fold-change in receptor expression at steady state and is discussed further below. *T*_fold_ is derived in Appendix A.1. To our knowledge, it has not been established yet how much less than one the Thiele modulus must be for the assumption of tissue homogeneity to hold.

### AFTIR derivation

#### AFTIR definition

Average Free Tissue target to Initial target Ratio (AFTIR) is defined as

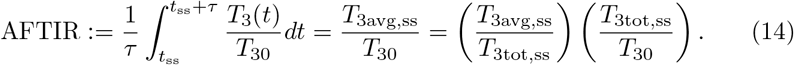

This equation applies for both membrane-bound (*M*) and soluble (*S*) targets, where *T*_3_(*t*) is the time series target concentration in the tumor compartment, *T*_30_ is the baseline target, *T*_3avg,ss_ is the average free target concentration at steady state, and *T*_3tot,ss_ denotes the total target concentration at steady state. AFTIR gives a measure of target inhibition in the tissue of interest (compartment 3) where the lower the AFTIR value, the greater the inhibition. The objective of this section is to derive the following estimate for AFTIR:

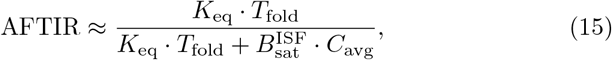

where *K*_eq_ is an equilibrium constant measuring how fast the drug and target bind to form complexs, *T*_fold_ is the fold-change in the target levels at steady state compared to baseline, 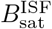 is the biodistribution coefficient (i.e. the fraction of the drug from circulation that is found in the tissue of interest), and *C*_avg_ is the average drug concentration in circulation. Expressions for each of these terms are derived below.

#### K_eq_ definition

Three different equilibrium constants are defined below. (1) *The quasi-equilibrium approximation* for both membrane and soluble target is [8]

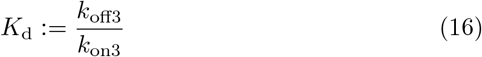

(2) *The quasi-steady state approximation* for soluble targets is [8]

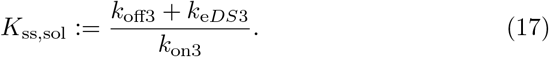

For membrane-bound targets, the formula is similar, though shedding needs to be taken into account as well:

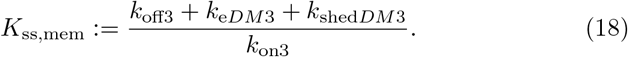

(3) A new equilibrium constant is introduced in this paper, called the *quasisteady-state with distribution constant*, denoted *K*_ssd_. This new equilibrium constant outperforms *K*_d_ and *K*_ss_ in terms of accuracy of approximating AFTIR.

The derivation of *K*_ssd_ for membrane-bound targets is shown below, and a similar process is followed for the soluble target version. To begin, assume a constant infusion of drug and that the system has reached steady state. Setting Equation (10) to zero and rearranging terms gives

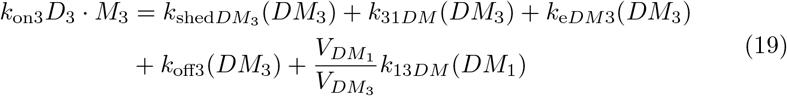

Dividing each side by *k*_on3_ · *DM*_3_ gives

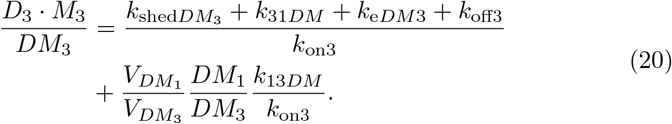

We then define *K*_ssd_ (for membrane-bound target and soluble target) as

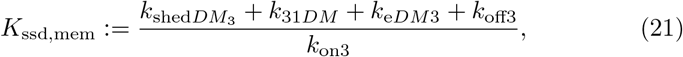

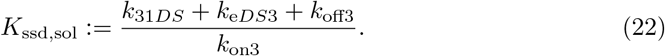

Substituting Equation (21) into Equation (20) gives

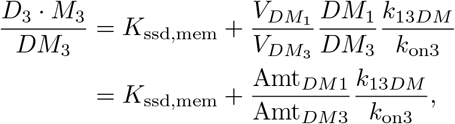

where 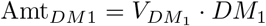 is the amount of complex in the central compartment, and 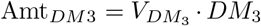 is the amount of complex in the tissue compartment. If the membrane-bound target is unable to move to the central compartment (i.e., *k*_13*DM*_ = *k*_31*DM*_ = 0), or if the amount of complex is much larger in the tissue than in circulation such that Amt_*DM*1_/Amt_*DM*3_ ≈ 0, then

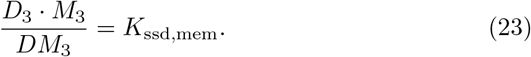

And similarly for soluble targets,

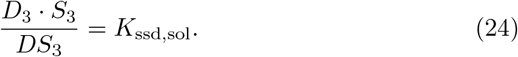

Regardless of which *K*_eq_ is used, the following approximation is assumed to hold:

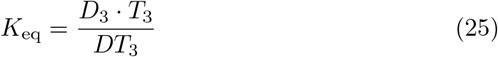

#### T_fold_, 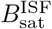, and C_avg_ definition

The total target in tissue is defined as *T*_3tot_ := *T*_3_+*DT*_3_. The fold-accumulation of the target in tissue at steady state is defined as

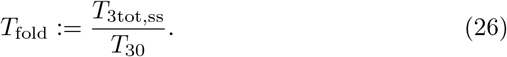

The analytical formula for *T*_fold_ is derived in Appendix (A.1) for the scenario where there is shedding of membrane-bound target to form soluble target. The scenario where no shedding takes place is calculated by setting either all shedding rates (*k*_shed_) terms to zero (so that there is no shedding), or, for soluble targets, by setting all membrane-bound target synthesis rates (*k*_syn*M*_) to zero.

For large drug concentrations that saturate the target, the steady state biodistribution coefficient 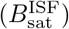 gives the the ratio of the drug concentration in tissue interstitial fluid (ISF) to that in circulation. The average steady state circulating concentration is defined as *C*_avg_ := *D*_1avg,ss_. The analytical formula for 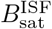 is derived in Appendix (A.2) and is in Equation 27.

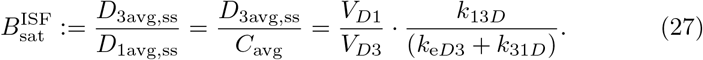

The above formula only applies to large doses that saturate the target. For low doses that do not saturate the target, binding to the target can significantly impact amount of drug that distributes to the tissue. Besides 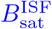, there are two other biodistribution terms that are also referenced in the literature: *B*_free_(*C*_avg_, *T*, *K*_eq_) and *B*_tot_(*C*_avg_, *T*, *K*_eq_), which refer to the fraction of free drug and total drug that is found in tissue compared to circulation. Both *B*_free_ and *B*_tot_ depend on the circulating drug concentration, level of target expression, and binding affinity. Throughout this manuscript, it is 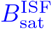 that will be used.

#### Putting it all together

Using the definitions of *T*_3tot_ and *K*_eq_, substituting the equation *DT*_3_ = *T*_3tot_ − *T*_3_ into the equation *K*_eq_ = *D* · *T*/(*DT*), and solving for *T*_3_/*T*_3tot_ gives

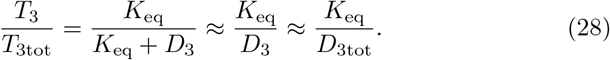

The first approximation holds when *D*_3_ ≫ *K*_eq_, and the second approximation holds when *D*_3tot_ ≈ *D*_3_, which occurs when drug is dosed in vast molar excess to the target, as is the case for most monoclonal antibody (mAb) drugs at the approved dosing regimen.

Using Equations (26) and (28) gives

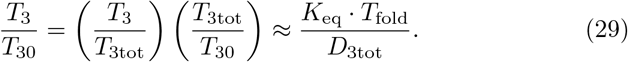

Integrating this ratio over one dosing cycle at steady state gives

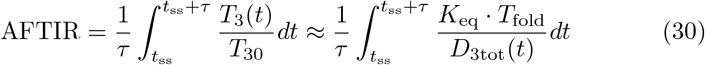

When the drug is given as an infusion at a rate Dose/*τ*, then the steady-state drug concentration is a constant (*D*_3tot,ss_), giving

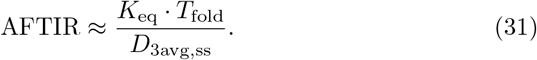

Substituting the biodistribution expression from Equation (27) gives what we call a simple expression for AFTIR.

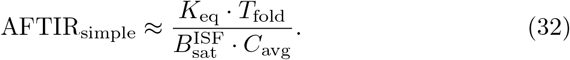

#### TEC_50_ definition

In deriving the above formula for the AFTIR metric, which we call the “simple” AFTIR formula, large drug concentrations are assumed. However, for arbitrarily small drug concentrations, AFTIR_simple_ is unbounded:

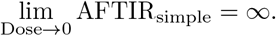

Realistically, AFTIR should approach one for arbitrarily small drug concentrations since the average free tissue target at steady state should approach the baseline tissue target concentration. Motivated by [9], we then modify the AFTIR approximation by adding an additional factor to the denominator:

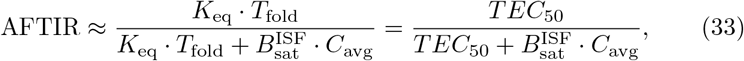

where the approximate concentration for 50% target engagement (*TEC*_50_) is given by

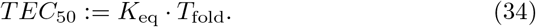

*TEC*_50_ is equivalent to the “L50” parameter in [9]. With this reformulation, we have

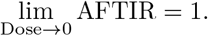

This new formulation still converges to the previously derived formula such that

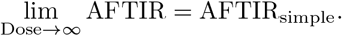

Both the *TEC*_50_ and simple formulas for AFTIR are checked against numerical simulations in the Results section. We attempted to derive the above expressions for AFTIR by extending the work from [9], but the derivation proved difficult for the model shown here. Thus, we instead posit that this approximation holds and will check it in the following sections.

#### Definition of TFTIR

Besides AFTIR, another quantity that can be used to guide dosing regimen is TFTIR (the Trough Free Tissue target to Initial target Ratio). It measures the minimum ratio of the free tissue target (e.g. within a tumor) to the initial tissue target at steady state and is defined as

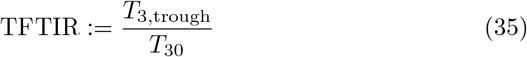

Like AFTIR, TFTIR can be approximated by

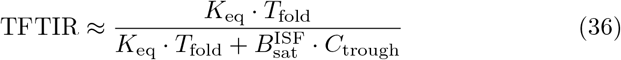

From [10], *C*_trough_ is given by

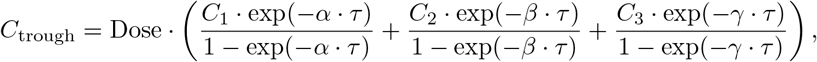

where all constants can be computed from the model parameters [11]. We will numerically check whether this approximation for TFTIR is accurate in the next section.

#### Linear PK and CL

In the case that the PK is linear, and there is no elimination from the tumor compartment (i.e., *k*_e*D*3_ = 0), one can write *C*_avg_ in terms of the dosing regimen and the parameters governing the pharmacokinetics: bioavailability (*F*), clearance (*CL*), and dosing interval (*τ*). It is given by

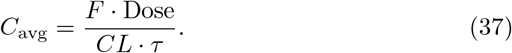

If there is elimination from the tumor (i.e., *k*_e*D*3_ ≠ 0), then the formula above does not apply with *CL* = *k*_e*D*1_/*V*_*D*1_. However, a similarity transform can be applied where *k*_e*D*3_ can be forced to be zero, and all the other PK parameters *k*_12*D*_, *k*_21*D*_, *k*_13*D*_, *k*_31*D*_, *k*_e*D*1_, *CL* can be transformed such that the concentration time profiles for *D*_1_(*t*) and *D*_3_(*t*) do not change. See Appendix A.3 for this similarity transformation. Thus, once *CL* is estimated by fitting a compartmental model, the formula above for *C*_avg_ holds whether or not there is elimination from the tissue compartment.

## Methods

Under the assumption of large doses and a constant infusion of drug, we have arrived at the following expression for AFTIR:

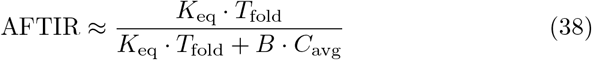

We hypothesize that this expression also holds for TFTIR, replacing *C*_avg_ with *C*_trough_.

To check the accuracy of this formula, we compare the theoretical AFTIR to the simulated AFTIR. The theoretical AFTIR is computed by Equation (38) for a given set of parameters. The simulated AFTIR is computed directly by simulating Equations (2) – (12) for the same set of parameters and then taking the ratio of the steady state average (or trough) free target concentration to the baseline target concentration. In addition, the Thiele Modulus is also computed using Equation (13). The model is implemented using the RxODE package in R, and correct implementation of the model is checked by comparing the simulated and analytical expressions for 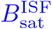, *C*_avg_, *C*_trough_ and *T*_fold_.

This simulations are performed for four mAbs that are used to treat cancer: trastuzumab (anti-HER2), atezolizumab (anti-PD-L1), pembrolizumab (anti-PD-1), and bevacizumab (anti-VEGF). Key properties of these drugs are summarized in Table (1), and the specific parameters used in this analysis are provided in Table S1. The references and equations used for selecting each parameter are provided in the Excel spreadsheets available here: https://github.com/iamstein/TumorModeling/tree/master/data and in the Supplementary Material. An overview for how the parameters are selected is provided below. For atezolizumab and pembrolizumab, membrane-bound target is shed to form soluble target. For trastuzumab, only membrane-bound target is simulated (shedding is set to zero), while for bevacizumab, only soluble target is simulated (synthesis of membrane-bound target is set to zero).

**Table 1.** Summary of drugs in this analysis. Details on all model parameters are provided in the Supplementary Material.

| Drug | Bevacizumab | Pembrolizumab | Atezolizumab | Trastuzumab |
| --- | --- | --- | --- | --- |
| Target Type | Soluble | Membrane-Bound | Membrane-Bound | Membrane-Bound |
| Target | VEGF | PD-1 | PD-L1 | HER-2 |
| Baseline Target (nM) | 0.01 | 0.2 | 3 | 800 |
| Shedding Included | no | yes | yes | no |

### Parameter Selection

The PK parameters (*k*_e*D*1_, *k*_12*D*_, *k*_21*D*_, *V*_*D*1_, *V*_*D*2_) are taken from PopPK model fits from the literature. The volumes for the target tissue are assumed to be 0.1 L, corresponding to a tumor with radius of 3 cm. The binding affinity (*K*_d_ or *K*_ss_) is also taken from the literature, and, when needed, a typical value for *k*_off3_ between 1/day-100/day was assumed [12, Figure 12]. The binding and unbinding rates are assumed to be the same in both the central and tissue compartments. For baseline soluble and membrane-bound target levels, data from the literature is used. For membrane-bound targets, target expression (*M*_10_, *M*_30_) is provided in terms of receptors per cell (*N*), which is then converted to nanomolar concentration with

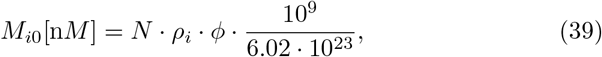

where *i* = 1, 3 refer to the central and target tissue compartments respectively. Here *ρ*_*i*_ is the cell density of the tissue of interest (*ρ*_1_ = 6·10^9^ cells/L in blood for targets expressed on white blood cells, and *ρ*_3_ = 3·10^11^ cells/L for targets expressed in tumor tissue [7]), and *ϕ* is the fraction of cells in the tissue of interest expressing the target, assumed to be 0.1 − 1.

The target elimination and shedding rates are chosen to have reasonable values (1/day – 10/day) [13], and the synthesis rates are chosen so that baseline target levels matched the estimates for *M*_*i*0_ above.

It is assumed that the rates of distribution of drug into the tumor are proportional to the rates of distribution into the peripheral tissue, scaled by the ratio of tissue volumes:

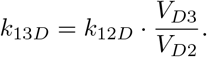

These estimates were similar to estimates from a more physiological model that accounted for drug perfusion into the tumor [7], as documented in the parameter Excel tables. The biodistribution coefficient 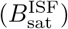 is assumed to be around 30% [4], and assuming no elimination of drug from the tumor, *k*_31*D*_ is estimated by assuming that at steady state, the transit rates into and out of the tumor are equal:

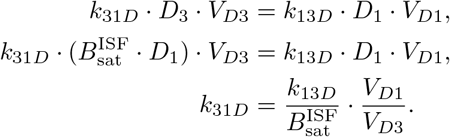

Soluble targets are then assumed to distribute two times faster than the drug, and soluble drug-taget complexes are assumed to distribute two times slower than the drug. It is recognized that this approach is empirical, and as a check, these rates were found to be comparable to the rates estimated using the Krogh cylinder approach for modeling tissue distribution [14].

For membrane-bound targets, where immune trafficking may occur (pembrolizumab, atezolizumab), it is assumed that the rate of distribution of cells expressing those targets into and out of tissue is *k*_13*M*_ = 0.048*/d*, and *k*_31*M*_ = 5.5*/d* [15, Table 4, parameters *k*_in_ and *k*_out_].

### Further parameter exploration

In the above definition of AFTIR, three different approximations are used for the equilibrium binding constant *K*_eq_: quasi-equilibrium (*K*_d_), quasi-steady-state (*K*_ss_), and the newly derived constant, quasi-steady-state with distribution (*K*_ssd_). The accuracy of each of these constants over a range of doses is compared for the four drugs above.

To further check the accuracy of the AFTIR approximation, we varied a few of the 47 model parameters over a large range with the dose fixed at a dose of 10 mg/kg every three weeks.

– *k*_13*D*_ – the rate of distribution of the drug from the central compartment to the tissue compartment. This will impact the biodistribution 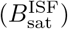 of drug in the tissue.
– *k*_13*DT*_ – the rate of distribution/trafficking of the drug-target complex from the central compartment to the tissue compartment. This parameter is included because it is often excluded from TMDD models and it was found to impact AFTIR.
– *k*_syn*T*3_ – the rate of target synthesis in the tissue compartment. This will impact whether the drug at a particular dose is in excess to its target.
– *k*_shed*DM*3_ – the rate of shedding of the drug-target complex from membrane-bound to soluble. This parameter is included because it is also often excluded from TMDD models.

For pembrolizumab, atezolizumab, and trastuzumab, *T* denotes the membrane-bound target, and for bevacizumab, *T* denotes the soluble target.

To determine the importance of an accurate estimate for the soluble target accumulation in the tissue of interest, we also explore how AFTIR varies for the parameter *k*_*eS*3_ for bevacizumab, as this parameter has a significant impact on *T*_fold_ = *S*_3tot,ss_*/S*_30_.

All code for generating the figures shown here is provided at: https://github.com/iamstein/TumorModeling/tree/master/ModelF.

## Results

AFTIR calculated by simulation is compared to the theoretical formula in Equation (38). The results are shown in Figures (2, 3, and 4).

**Fig. 2.**
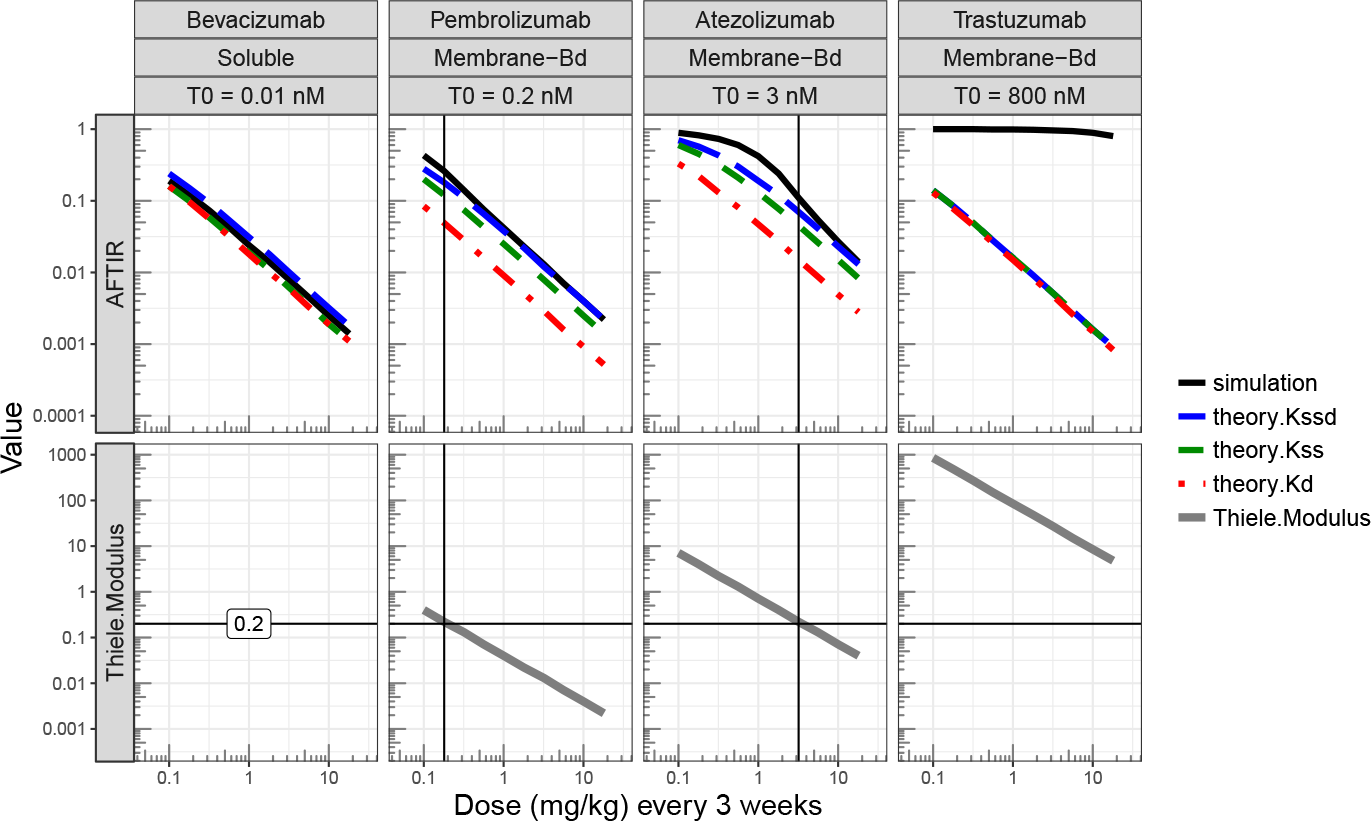
Average Free Tissue target to Initial target Ratio (AFTIR, top row) and Thiele Modulus (*ϕ*^2^, bottom row) for four drugs with different baseline target expression levels (*T*_0_) over a range of doses. The *K*_ssd_ approximation matches the simulation better than *K*_ss_ and *K*_d_. When the Thiele Modulus is less than 0.2, the AFTIR *K*_ssd_ approximation matches the data well. For trastuzumab in particular, *ϕ*^2^ > 10, target expression is high (800 nM), and so the theory does not match the simulation.

**Fig. 3.**
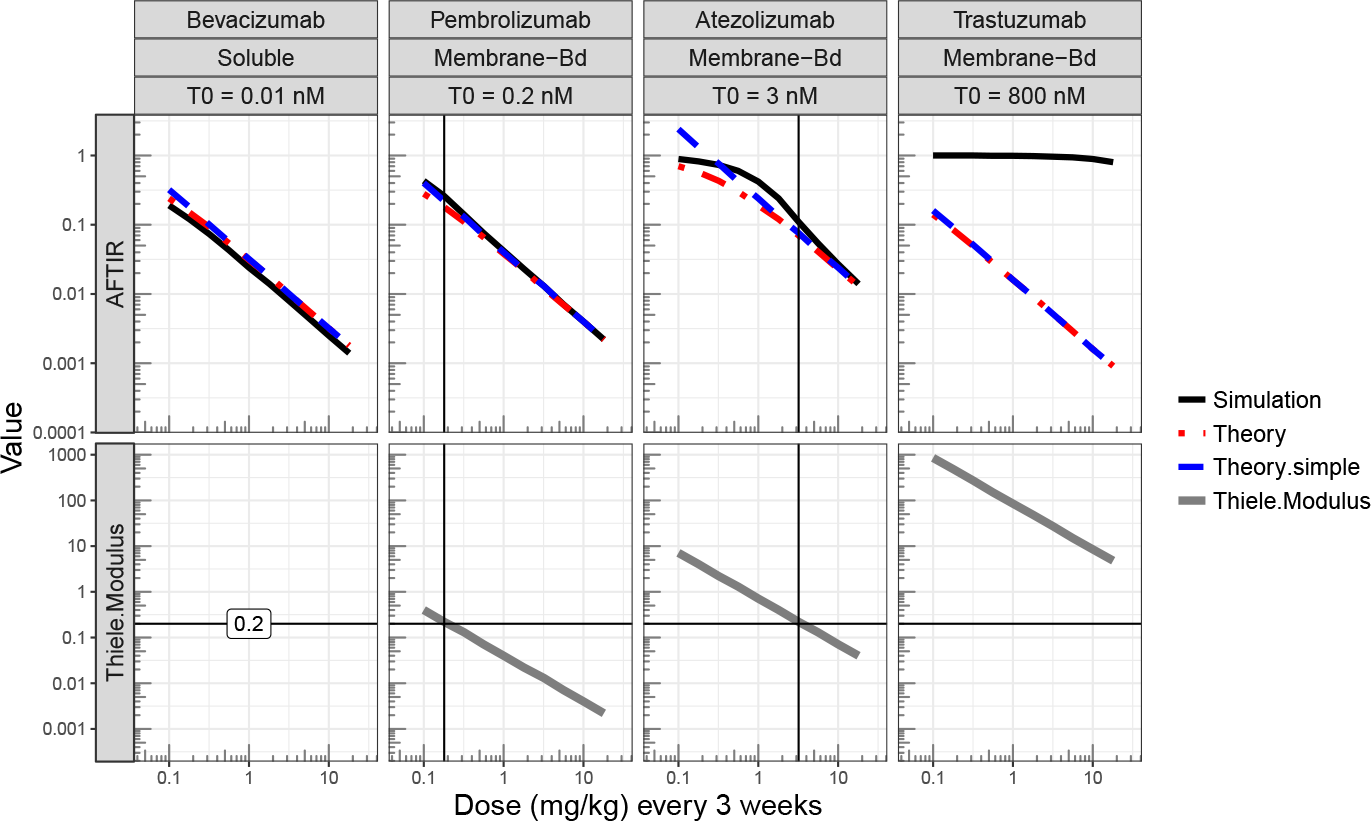
Average Free Tissue target to Initial target Ratio (AFTIR, top row) and Thiele Modulus (*ϕ*^2^, bottom row) for four drugs with different baseline target expression levels (*T*_0_) over a range of doses. The AFTIR approximation better matches the simulation than the AFTIR_simple_ approximation for very small doses because at these doses, AFTIR asymptotically approaches 1, whereas AFTIR_simple_ is unbounded.

**Fig. 4.**
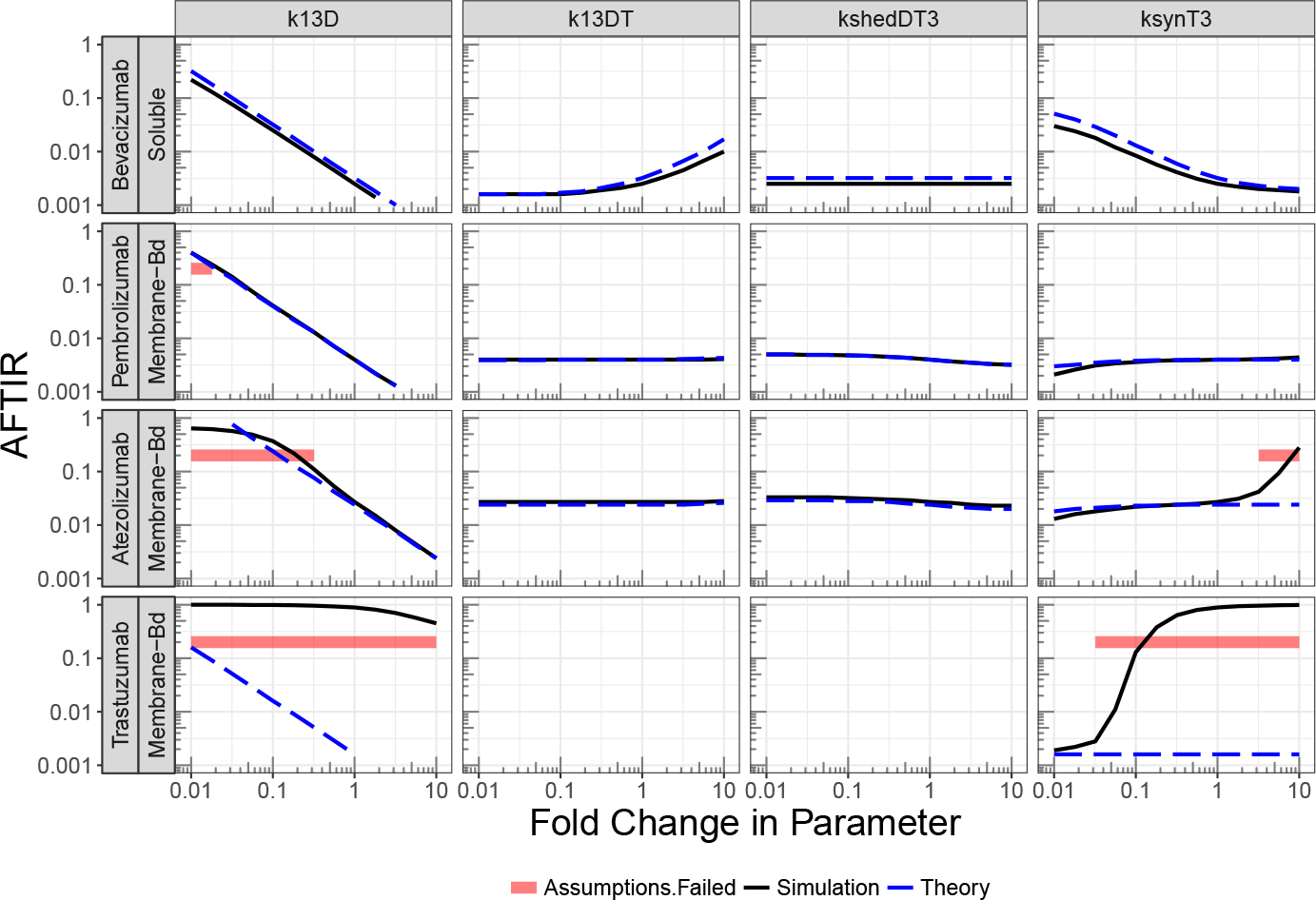
Exploration of how other parameters impact the Average Free Tissue target to Initial target Ratio (AFTIR) using the *K*_ssd_ approximation. The parameter at the top of each column is changed from 0.01× to 10× from its baseline value. The red line indicates the regime where the assumption of tissue homogeneity does not hold. Here, this assumption is checked by testing whether the Thiele Modulus is greater than 0.2. It is in the regime covered by the red line that the theory and simulation do not agree.

Figure (2) (top row) shows comparisons of the approximations over a range of doses using the three equilibrium constants, *K*_d_, *K*_ss_, and *K*_ssd_. In general, *K*_ssd_, the new equilibrium constant derived here, best matches the model simulation. Note, however, that for trastuzumab, the drug with the highest target levels, the theory does not match the simulation at all. This is because the AFTIR approximation requires the assumption that the drug is in vast excess to its target and that the tumor can be treated as a homogeneous tissue. The bottom row of Figure (2) shows how the Thiele modulus changes with dose. When the Thiele Modulus is less than 0.2, there is relatively good visual agreement between the theory and the simulation. Because the Thiele modulus is primarily applicable for membrane-bound targets, it is not computed for Bevacizumab.

Note that for Atezolizumab, the agreement improves when the dose drops to 0.1 mg/kg, despite the Thiele Modulus being greater than 0.2. This is because for any drug given at sufficiently low doses, *AFTIR* ≈ 1 for both theory and simulation because no drug is present.

Another feature to note is that the *K*_ss_ approximation differs from the *K*_ssd_ approximation by less that 2-fold. Thus, the *K*_ss_ approximation is still reasonable, and if one wanted a more conservative estimate for target engagement, a 2× “safety factor” could multiply *K*_ss_. Similar findings are also observed for TFTIR (see supplementary material).

In Figure (3), it is shown that at high doses, AFTIR and AFTIR_simple_ agree well at values below 0.3. But above 0.3, and especially at small concentrations where AFTIR approaches 1, the simple approximation no longer holds, as expected.

Figure (4) shows the sensitivity of AFTIR to other parameters, namely the rate of trafficking of the drug from the central compartment to the tissue compartment (*k*_13*D*_), the rate of trafficking of the drug-target complex from the central compartment to the tissue compartment (*k*_13*DT*_), the rate of shedding of the drug-target complex from membrane-bound to soluble (*k*_shed*DM*3_), and the rate of target synthesis in the tissue compartment (*k*_syn*T*3_). The red horizontal line denotes the regime where the Thiele Modulus is greater than 0.2, indicating that the assumptions of tissue homogeneity and drug in excess of the target are inaccurate. For bevacizumab and pembrolizumab, there is generally good agreement between theory and simulation. For trastuzumab, the theory does not match the simulation due to the very high target concentration. And for atezolizumab, the theory matches the simulation only when the Thiele modulus is below 0.2

For atezolizumab, the parameter for which there is the greatest discrepancy between theory and simulation is *k*_syn*T*3_. At low *k*_syn*T*3_ values, the theory overpredicts the AFTIR value, while for high *k*_syn*T*3_ values, the theory under-predicts the AFTIR value. The reason for the discrepancy at high *k*_syn*T*3_ values is that the target expression becomes so high that the drug is no longer in excess to the target, as in the trastuzumab case. The reason for the discrepancy at low *k*_syn*T*3_ values is that the assumption of negligible complex transport from circulation into the tissue of interest no longer applies. And so the approximation for deriving *K*_ssd_ in Equation (21) is no longer accurate. As a check of this phenomenon, we set *k*_13*DT*_ to zero. As a result, there is better agreement between theory and simulation in Figure (S1).

In Figure (5), it is shown that for a drug (bevacizumab) with a soluble target (VEGF), the fold-accumulation of the target is a critical factor in predicting the amount of target inhibition in the tissue of interest. This is also observed in Equation (31), where AFTIR is directly proportional to *T*_fold_.

**Fig. 5.**
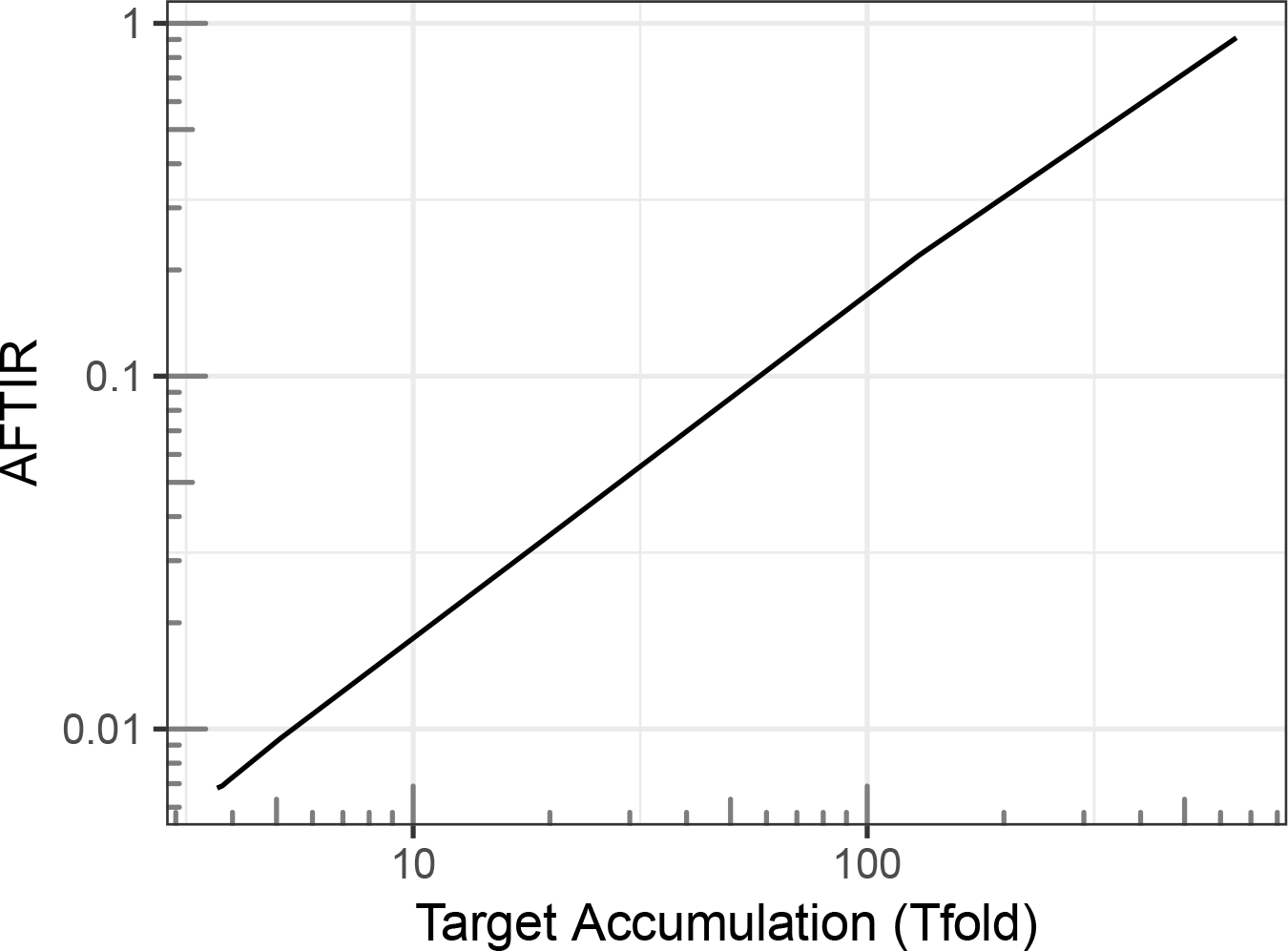
Simulated Average Free Tissue target to Initial target Ratio (AFTIR) as a function of fold change in target (*T*_fold_) for Bevacizumab.

## Discussion

### Review of key AFTIR parameters

The key insight from this work is that under many clinically relevant scenarios, AFTIR can be estimated using four parameters, a binding constant (*K*_eq_), the fold-change in target levels upon binding to drug (*T*_fold_), the average drug concentration in circulation (*C*_avg_), and the biodistribution coefficient for the drug to the tissue of interest 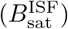:

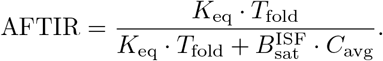

This simple formula provides intuition for how changing the dosing regimen, improving the binding affinity of the drug, or enhancing tissue penetration would be expected to alter target inhibition. For example, at large drug concentrations, when 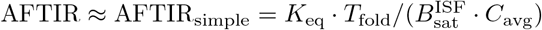, one can see that either halving the dosing interval, halving the binding affinity (with a more potent drug), or doubling the tissue accessibility (with a drug that enhances tissue penetration) would reduce the free target concentration by approximately 50%. Even though the model in Figure (1) has many parameters, some of which are difficult to estimate, the AFTIR formula shows that under many practical scenarios, predicting the target inhibition can be done with an estimate of only four lumped parameters. The method for estimating each of these parameters is described below.

The **drug concentration** (*C*_**avg**_ **or** *C*_**trough**_) is estimable from PK data from Phase 1 clinical trials. For monoclonal antibodies, it can also be readily predicted from preclinical data [16].

The **binding affinity** (*K*_**d**_,*K*_**ss**_ **or** *K*_**ssd**_) can either be estimated in vitro with surface plasmon resonance and cell binding assays, or in the clinic with a model-based analysis of soluble target data [1] or receptor occupancy assays [17]. Often there is good agreement between the in vitro and in vivo estimates, though it is critical that the assays be carefully validated as there have been instances of 1000-fold differences between the in vitro and in vivo estimates [18, Figure 8]. Also, assays measuring free or bound target concentrations, rather than total target concentrations, can be difficult to validate [19]. While *K*_ssd_ provided the best match between theory and simulation, it is difficult to measure directly because it requires target occupancy measurements in the tissue of interest. Thus a modeler may also use an estimate for *K*_ss_ and, if desired, multiply this binding affinity by a safety factor (e.g. 2×) for a more conservative estimate of target engagement.

The **biodistribution coefficient** 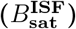 has been estimated for many tissues in monkeys [20], where it ranges from 5 − 15% in the total tissue or 15 − 45% in the tissue interstitial fluid (assuming that 1/3 of the tissue is interstitial fluid). In mouse tumors, 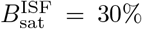 [4], and this is similar in human skin for secukinumab [21]. However, others have used predictions of tumor distribution based on rates of extravasation and diffusion to predict that the antibody concentration in a tumor should be only 0.1−1% [6]. Due to this disagreement in estimates for 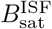 and the limited data in the literature, further work in estimating biodistribution would be of value [22].

The **fold-change in target concentration** (*T*_**fold**_) in the tissue of interest can be estimated from in vitro experiments for membrane-bound targets by estimating the internalization rates of bound and unbound receptors. It is generally around *T*_fold_ = 0.5 [23], [24, page 15]. Care must be taken when interpreting biopsy staining data from immunohistochemistry (IHC) assays to estimate *T*_fold_.

For soluble targets, *T*_fold_ is often directly measured in circulation [1], but to our knowledge, it has only been measured once in tissue, namely for IL-6 in mouse synovial fluid [25]. This data was difficult to interpret since many measurements were below the limit of quantification of the assay. Therefore, a significant challenge in applying this formula to soluble targets is that the extent of target-accumulation in the tissue of interest is not known. Some of the methods for estimating 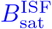 can also be employed for measuring *T*_fold_; in particular, techniques such as dermal open flow microperfusion (skin) [21] or microdialysis (tumor) [26] could be employed to measure both drug concentration and soluble target concentration.

### Review of assumptions and caveats

The following assumptions are needed for the theoretical estimate of AFTIR to be accurate.

#### The drug concentration is much larger than the target concentration, and the tissue can be treated as homogeneous

This assumption should be checked in two ways. First, by comparing estimates of target expression in the tissue of interest to estimates of the drug concentration in that tissue. And second, for membrane-bound targets, one can check whether the Thiele Modulus is small. [6, 7] For this model, it is found that the Thiele modulus has to be less than 0.2 for the theory to describe the simulation well. The assumption of homogeneity fails for trastuzumab at all doses and for atezolizumab at doses less than 3 mg/kg q3w. In the future, it would be useful to check whether this threshold of 0.2 holds for other drugs and other models.

#### Distribution of the drug-target complex from circulation into the tissue is relatively small

This assumption was required for the calculation of *K*_ssd_. If distribution of the complex from circulation into the tissue of interest plays a significant role, then the AFTIR formula may not apply. One instance where it may need to be checked is for a soluble target that accumulates 100–1000 times in circulation but not in the tissue of interest. This assumption is difficult to check and requires knowledge of the underlying biology. If it is known that the target is primarily expressed in the tumor, or if its a membrane-bound target that doesn’t move from circulation into the tumor, then this assumption would be reasonable.

#### The degree of inhibition needed for efficacy is known

Often, 90-95% inhibition is assumed to be needed [27], though for a drug that works by stimulating the immune system to attack the tumor, much less inhibition may be needed. Examples include trastuzumab, which may also work via antibody dependent cell-mediated cytotoxicity and bispecific T-cell engagers like blinatumomab, which facilitates the interaction between T cells and cancer cells.

#### Competition for target binding sites between drug and endogenous ligand

If the binding between a target and its endogenous ligand is much lower than the binding affinity of the drug, the formulas derived here may not applicable because the drug may not be disrupting the interactions between the target and its endogenous ligand. In this analysis, avidity is also not included in the analysis. In the scenario where the monoclonal antibody is in vast excess to the target, each drug molecule is expected to bind at most only one target molecule, and thus avidity was not addressed in this analysis.

#### Distribution of target and complex to peripheral tissue can be neglected

A more realistic model would include distribution of the drug, target, and complex to each tissue; a lumped, minimal PBPK model could also be considered that also allows for distribution of all components to peripheral tissue. However, we note that this approach using a peripheral compartment for just drug concentrations is often used for predicting tissue distribution [7, 28, 3]. Moreover, the expression for *T*_fold_ for this model was already quite complex, and thus the analysis of this more complex system is left to future research.

### Applications

#### Phase 2 dose selection

A key application of this work is to provide an updated methodology for guiding Phase 2 dose selection for biologics when the safety, efficacy, and biomarker data are insufficient to guide dose selection. With this approach, one uses a PopPK model based on Phase 1 PK data to predict the steady state trough concentration of the drug. Then, together with estimates for the binding affinity, fold accumulation of target upon treatment, and biodistribution of drug to tissue, one can apply the AFTIR or TFTIR equation to identify the dose at which, say, 90% of patients are expected to achieve 90% target inhibition. While a similar methodology was used previously for atezolizumab [4], *K*_ssd_ and *T*_fold_ were not explicitly accounted for.

#### Rapid assessment of drugs

The AFTIR potency metric also allows for a rapid assessment of new drugs without a need for extensive simulations. In particular, given that 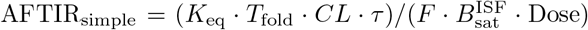, this formula shows how changing the dosing regimen, clearance, and binding affinity would impact target engagement.

Moreover, rather than specify every rate constant of the complex system of Figure (1), or an even more complex physiological model, it is sufficient in many scenarios to provide estimates for only four parameters: *K*_eq_, *T*_fold_, 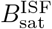, and *C*_avg_, all of which can in principle be measured directly.

#### Rapid sensitivity analyses

While all parameters can in principle be measured directly, there will often be scenarios where there is uncertainty about the specific parameter values. In that case, the AFTIR potency metric also allows for a rapid calculation of the impact of this uncertainty on the predicted receptor occupancy. Given the uncertainty, one can then either try to collect additional information about the parameters with the highest degree of uncertainty, or factor this uncertainty into the analysis when informing drug development decisions. For instance, if a drug is relatively safe, one may choose to take forward a higher dose into Phase 2 to ensure that more patients achieve a high degree of target engagement.

### Conclusions

In summary, we have extended previous work to develop potency metrics AFTIR and TFTIR that characterize target inhibition in a tissue, such as a tumor. These metrics predict the target inhibition at steady state under a repeated dosing regimen using four quantities: the average (or trough) drug concentration at steady state, the biodistribution of drug to its target tissue, the equilibrium binding constant, and the fold-accumulation of the target. AFTIR and TFTIR provide intuition for how various physiological properties of the system impact target engagement. In addition, AFTIR and TFTIR can be used for rapid assessment of new targets and exploration of the binding affinity, half-life, bioavailability, and dosing regimen needed for a second-generation drug to achieve comparable or superior efficacy to a marketed compound. Thus far, a significant challenge in applying AFTIR is in obtaining accurate, clinical estimates in tissue for biodistribution and fold-accumulation of target in the tissue of interest. It is our hope that this work will motivate scientists to use assays like dermal open flow microperfusion or microdialysis to better estimate these parameters in the future.

## Acknowledgements

The authors would like to thank the IMA as the Math-to-Industry Bootcamp was where this work began. We’d also like to thank Ngartelbaye Guerngar who was also a member of the bootcamp team and Wenping Wang and Matt Fidler for help in using the RxODE package. Finally, we’d like to thank Eduardo Sontag and Greg Thurber for many helpful discussions.

## Appendix

## Fold-Accumulation (*T*_fold_)

Recall that the fold-accumulation of target is given by

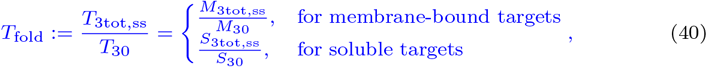

where *T*_3tot,ss_ denotes the sum of target and drug-target complex concentration at steady state, and *T*_30_ denotes target concentration at the initial state. In the sections below, we solve for the baseline target and the steady state target (for large drug concentrations) using similar approaches. Then, we compute *T*_fold_ as the ratio of the two.

## Baseline target (*T*_10_ and *T*_30_)

Before the drug is given, the system is at steady state. And we have *D*_1_ = *D*_2_ = *D*_3_ = *DM*_1_ = *DM*_3_ = *DS*_1_ = *DS*_3_ = 0. Using the ODE system, we solve for the nonzero initial membrane-bound targets. Setting Equations (5) and (6) to zero, we get a system of two linear equations,

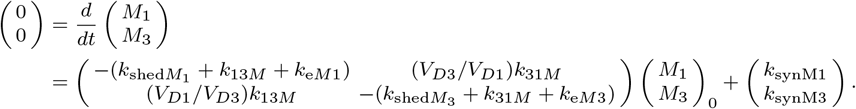

Rearranging yields

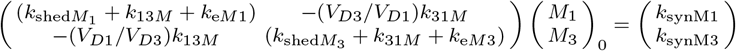

Then, using the formula for inverting a 2D matrix gives

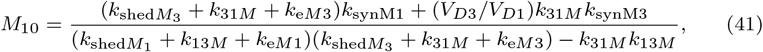

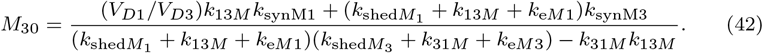

To solve for the nonzero initial soluble targets, we set Equations (7) and (8) to zero, resulting in a system of two linear equations,

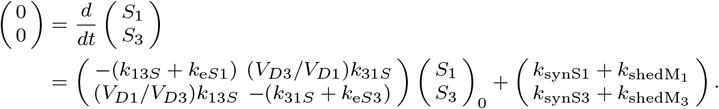

Similar to above, we get

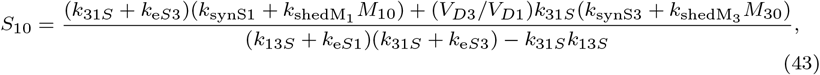

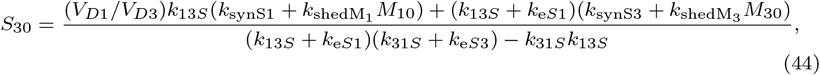

where *M*_10_ and *M*_30_ are given by (41) and (42).

## Steady State Value

After the drug is given, the system reaches steady state. We assume that the drug is in vast excess to the amount of target so that almost all of the target is bound to the drug. Then we have *M*_1_ = *M*_3_ = *S*_1_ = *S*_3_ ≈ 0, *M*_3tot,ss_ = *M*_3_ + *DM*_3_, and *S*_3tot,ss_ = *S*_3_ + *DS*_3_. Using the ODE system, we solve for the nonzero steady state of the total membrane-bound targets. Adding the equations for the target and complex, Equations (5) and (6) added to Equations (9) and (10), respectively, gives differential equations for the total target that are similar to that when no drug is on board. We set these differential equations to zero, resulting in a system of two linear equations. We solve as in the previous section to get

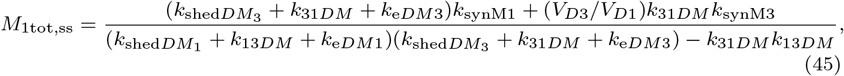

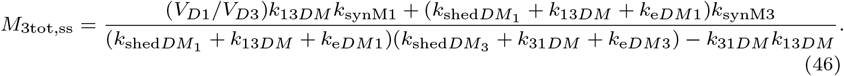

Likewise, the nonzero steady state of total soluble targets are given by

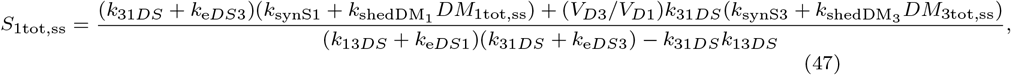

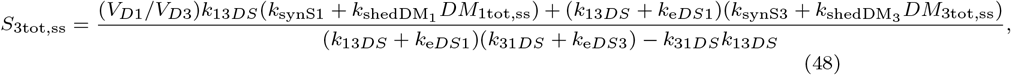

where *M*_1tot,ss_ and *M*_3tot,ss_ are given by (45) and (46).

A summary of all the of equations is provided below. To calculate *T*_fold_, one then uses the formulas below to compute *M*_3tot,ss_/*M*_30_ or *S*_3tot,ss_/*S*_30_.

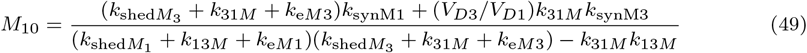

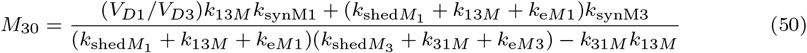

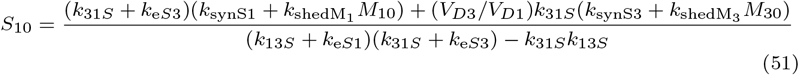

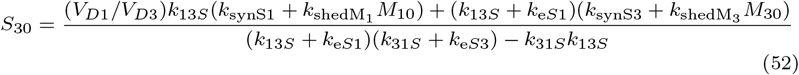

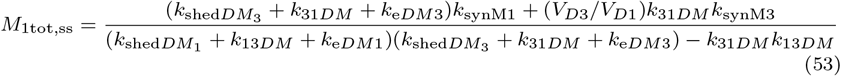

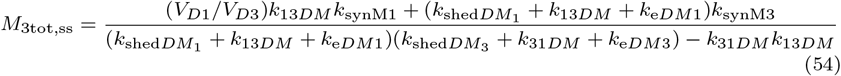

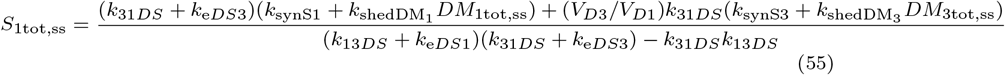

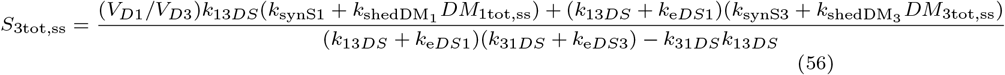

## Biodistribution 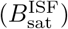

For very large infusion where the target concentration is negligable, we can write the three equations for the drug (*D*_1_, *D*_2_, *D*_3_) from Equations (2), (3), and (4) as below to solve for steady state.

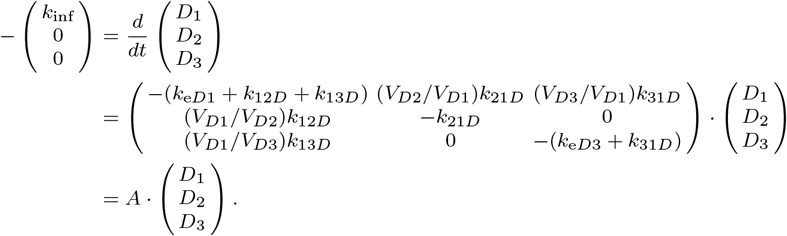

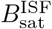 is given by the ratio of *D*_3,ss_/*D*_1,ss_ where *ss* denotes steady state, such that when the drug concentration is large

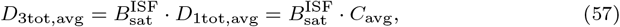

where we define *C*_avg_ := *D*_1tot,avg_.
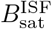 is computed as follows:

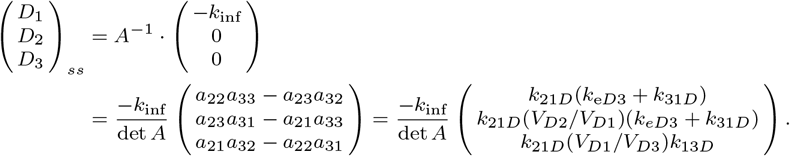

Thus the biodistribution coefficient for the tissue interstitial fluid compartment is given by

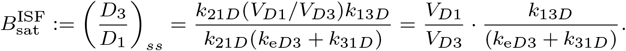

For drugs dosed with linear PK at regular intervals, drug concentration much larger than target concentration, and free drug elimination occuring only in the central compartment, we have *D*_1tot,avg_ = D_ose_/(*CL* · *τ*), where *CL* = *k*_e*D*1_ · *V*_*D*1_ is the drug clearance. As shown in Section A.3, we can derive a similar formula with coefficient matrix *A′* in which *k*_e*D*3_ = 0, while 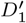 and 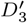 in the new model are the same as *D*_1_ and *D*_3_ in the original model. From Section A.3, we have

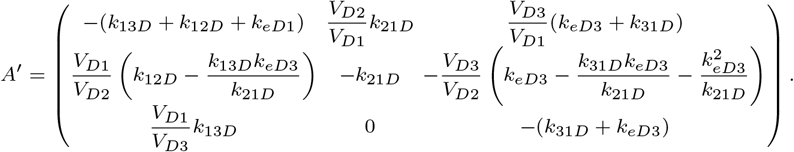

Then, as above,

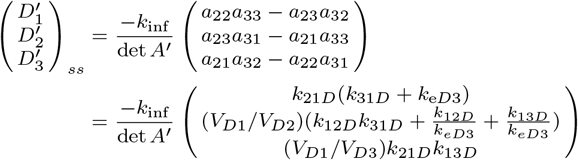

And the biodistribution coefficient for the tissue interstitial fluid compartment is given by

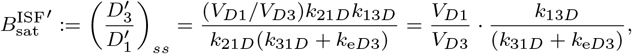

which is the same as 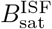 in the original model above.

## Similarity Transform

We consider the model in Figure (1). A drug (*D*) binds to its target (*M*) to form a complex (*DM*). It has three compartments, central, tissue, and peripheral.

The drug and target dynamics are modeled with the following system of ordinary differential equations

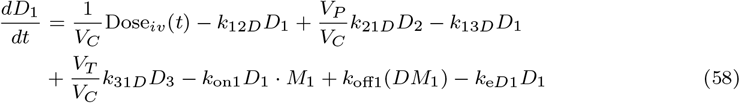

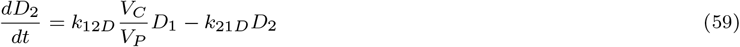

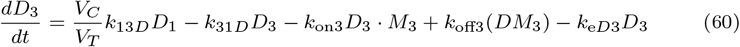

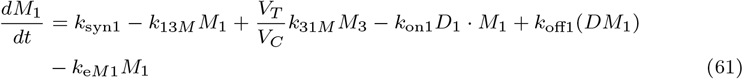

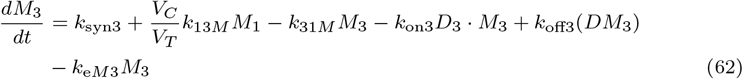

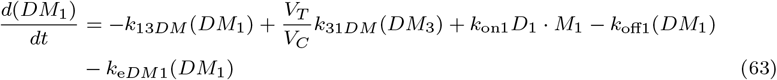

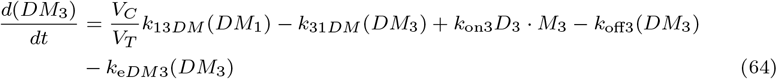

We will calculate *D*_3tot,avg_, which is the average drug concentration at steady state in the tissue compartment. We have 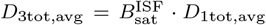, where *D*_1tot,avg_ is the average drug concentration at steady state in the central compartment, and 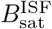 is the antibody biodistribution coefficient. Assume that drug elimination only occurs in the central compartment, i.e., *k*_e*D*3_ = 0. Then for drugs dosed with linear PK at regular intervals and concentration much larger than target concentration, we have *D*_1tot,avg_ = Dose/(*CL · τ*), where *CL* = *k*_e*D*1_ · *V*_*C*_ is the drug clearance [10].

For the model with drug elimination in both the central and tissue compartments, we will derive an alternative model with *k*_e*D*3_ = 0 that is indistinguishable from the existing model by using the similarity transform technique [29]. We consider drug concentration large enough that target binding does not affect drug distribution. Then the model equations for the drug can be written as

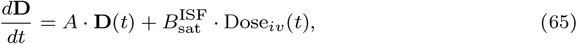

with measurement

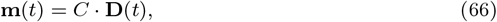

where

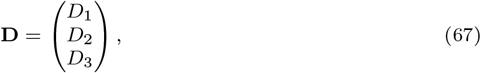

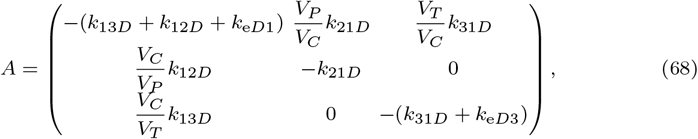

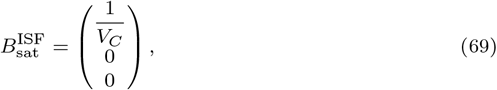

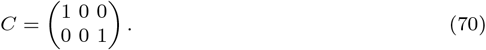

We transform this model into an indistinguishable model with coefficient matrices *A′*, 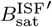, and *C′* such that 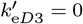. This is accomplished with the similarity transform

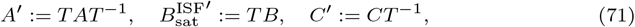

for some transformation matrix

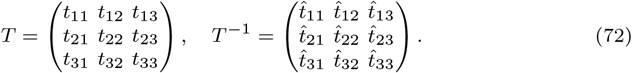

Alternatively, one can get this similarity transform with the change of variables **D** = *T*^−1^**D*′***. Then from (65) and (66), we have

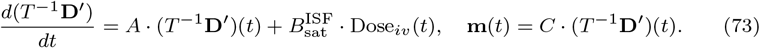

Multiplying the first equation in (73) on the left by *T* yields

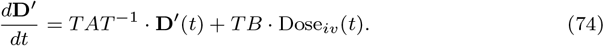

Then we have the new model equation and measurement

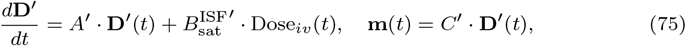

where *A′*, 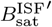, and *C′* are given by (71).

Regardless of model formulation, the same dose is given to *D*_1_, that is,

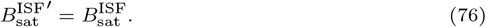

From this and (71), we get *t*_11_ = 1, *t*_21_ = 0, and *t*_31_ = 0. Regardless of model formulation, we measure *D*_1_ and *D*_3_, that is,

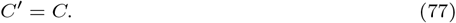

From this and (71), we get 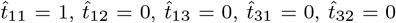, and 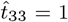. For a 3 × 3 matrix, the inverse is given by

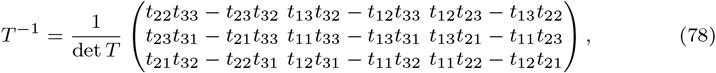

with the determinant given by

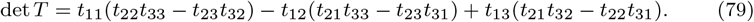

Putting the above findings into *T*^−1^ yields *t*_12_ = 0, *t*_13_ = 0, *t*_32_ = 0, and *t*_33_ = 1. From *TT*^−1^ = *I*, we get *t*_22_ = 1. Then we have

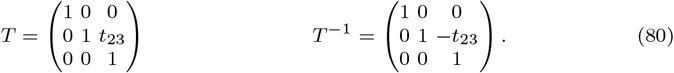

Putting this into (71) yields

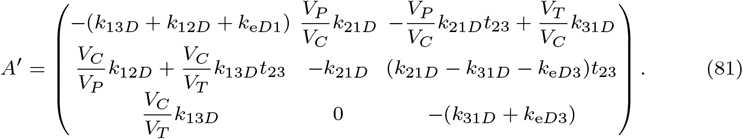

Now we impose

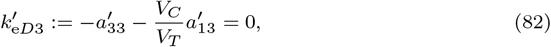

from which we get *t*_23_ = − (*V*_*T*_ /*V*_*P*_)(*k*_e*D*3_/*k*_21*D*_). After putting this last piece into *A′*, we have

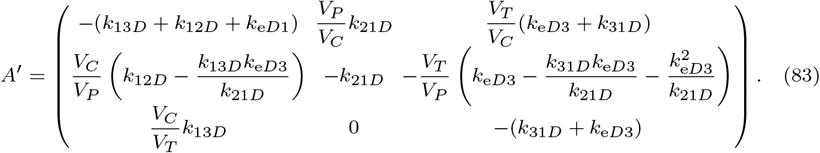

We get 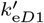 from

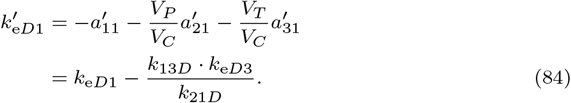

Thus 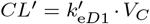, with 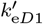 given by (84). And finally we get *D*_tot,avg1_ = Dose/(*CL′ · τ*), and 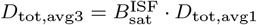.

*Remark* : In the similarity transform, we measure *D*_1_ and *D*_3_, which is indicated by *C*. As a result of imposing *C′* = *C*, the original model and the transformed model output the same values for these variables. 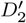, however, differs from *D*_2_.

## Supplementary Material

All parameters and code for executing the model are stored here: https://github.com/iamstein/TumorModeling

### Model Parameters

**Table S1.** Parameters used for the simulations. To find the references and calculations for how these parameters were derived, consult https://github.com/iamstein/TumorModeling/tree/master/data

| Parameter | Units | Atezolizumab | Bevacizumab | Pembrolizumab | Trastuzumab |
| --- | --- | --- | --- | --- | --- |
| $k_{12D}$ | 1/d | 0.17 | 0.43 | 0.23 | 0.17 |
| $k_{13D}$ | 1/d | 0.0046 | 0.014 | 0.0056 | 0.013 |
| $k_{13DM}$ | 1/d | 0.048 | 0 | 0.048 | 0 |
| $k_{13DS}$ | 1/d | 0.0046 | 0.014 | 0.0056 | 0 |
| $k_{13M}$ | 1/d | 0.048 | 0 | 0.048 | 0 |
| $k_{13S}$ | 1/d | 0.0092 | 0.028 | 0.011 | 0 |
| $k_{21D}$ | 1/d | 0.15 | 0.45 | 0.2 | 0.28 |
| $k_{31D}$ | 1/d | 0.45 | 1.3 | 0.59 | 0.83 |
| $k_{31DM}$ | 1/d | 5.5 | 0 | 5.5 | 0 |
| $k_{31DS}$ | 1/d | 0.45 | 1.3 | 0.59 | 1 |
| $k_{31M}$ | 1/d | 5.5 | 0 | 5.5 | 0 |
| $k_{31S}$ | 1/d | 0.9 | 2.7 | 1.2 | 1 |
| $k_{eD1}$ | 1/d | 0.061 | 0.056 | 0.063 | 0.1 |
| $k_{eD3}$ | 1/d | 0 | 0 | 0 | 0 |
| $k_{eDM1}$ | 1/d | 0 | 1 | 0 | 1 |
| $k_{eDM3}$ | 1/d | 0 | 1 | 0 | 2.9 |
| $k_{eDS1}$ | 1/d | 0.005 | 0.07 | 0.063 | 1 |
| $k_{eDS3}$ | 1/d | 0 | 0.07 | 0 | 1 |
| $k_{eM1}$ | 1/d | 0 | 1 | 0 | 1 |
| $k_{eM3}$ | 1/d | 0 | 1 | 0 | 2.9 |
| $k_{eS1}$ | 1/d | 0.3 | 7 | 0.47 | 1 |
| $k_{eS3}$ | 1/d | 0 | 7 | 0 | 1 |
| $k_{off1}$ | 1/d | 3 | 2 | 3.5 | 30 |
| $k_{off3}$ | 1/d | 3 | 2 | 3.5 | 30 |
| $k_{on1}$ | 1/(nM*d) | 1.2 | 36 | 8.2 | 60 |
| $k_{on3}$ | 1/(nM*d) | 1.2 | 36 | 8.2 | 60 |
| $k_{shedDM1}$ | 1/d | 6 | 1 | 6 | 0 |
| $k_{shedDM3}$ | 1/d | 6 | 1 | 6 | 0 |
| $k_{shedM1}$ | 1/d | 3 | 1 | 3 | 0 |
| $k_{shedM3}$ | 1/d | 3 | 1 | 3 | 0 |
| $k_{synM1}$ | nM/d | 0.45 | 0 | 0.03 | 0 |
| $k_{synM3}$ | nM/d | 22 | 0 | 1.5 | 2400 |
| $k_{synS1}$ | nM/d | 0 | 0.014 | 0 | 0 |
| $k_{synS3}$ | nM/d | 0 | 0.1 | 0 | 0 |
| $V_{D1}$ | L | 3.3 | 3.2 | 3.5 | 2.1 |
| $V_{D2}$ | L | 3.6 | 3.1 | 4.1 | 1.3 |
| $V_{D3}$ | L | 0.1 | 0.1 | 0.1 | 0.1 |
| $V_{DM1}$ | L | 3.3 | 3.2 | 3.5 | 2.1 |
| $V_{DM3}$ | L | 0.1 | 0.1 | 0.1 | 0.1 |
| $V_{DS1}$ | L | 3.3 | 3.2 | 3.5 | 2.1 |
| $V_{DS3}$ | L | 0.1 | 0.1 | 0.1 | 0.1 |
| $V_{M1}$ | L | 3.3 | 3.2 | 3.5 | 2.1 |
| $V_{M3}$ | L | 0.1 | 0.1 | 0.1 | 0.1 |
| $V_{S1}$ | L | 3.3 | 3.2 | 3.5 | 2.1 |
| $V_{S3}$ | L | 0.1 | 0.1 | 0.1 | 0.1 |

### AFTIR Simulations

**Fig. S1.**
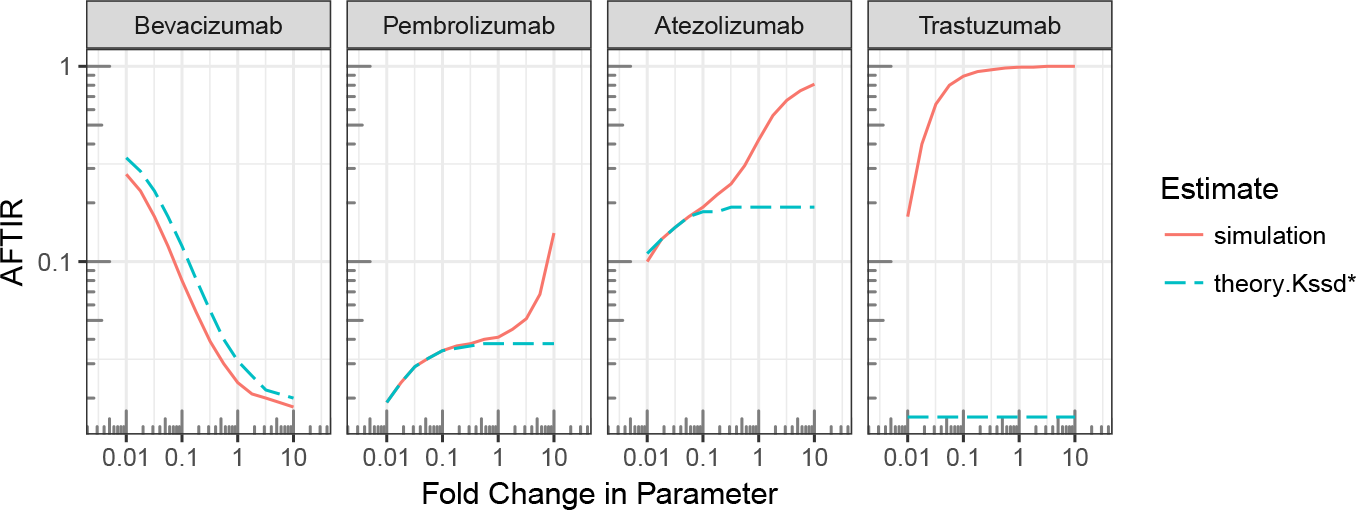
Parameter exploration for *k*_syn*T*3_. *k*_13*DT*_ has been set to zero gives better agreement for *k*_syn*T*3_. Here, the parameter at the top of each column is changed from 0.01× to 10× from its baseline value. There is in general good agreement between theory and experiment, except for trastuzumab, when the target expression is high.

**Fig. S2.**
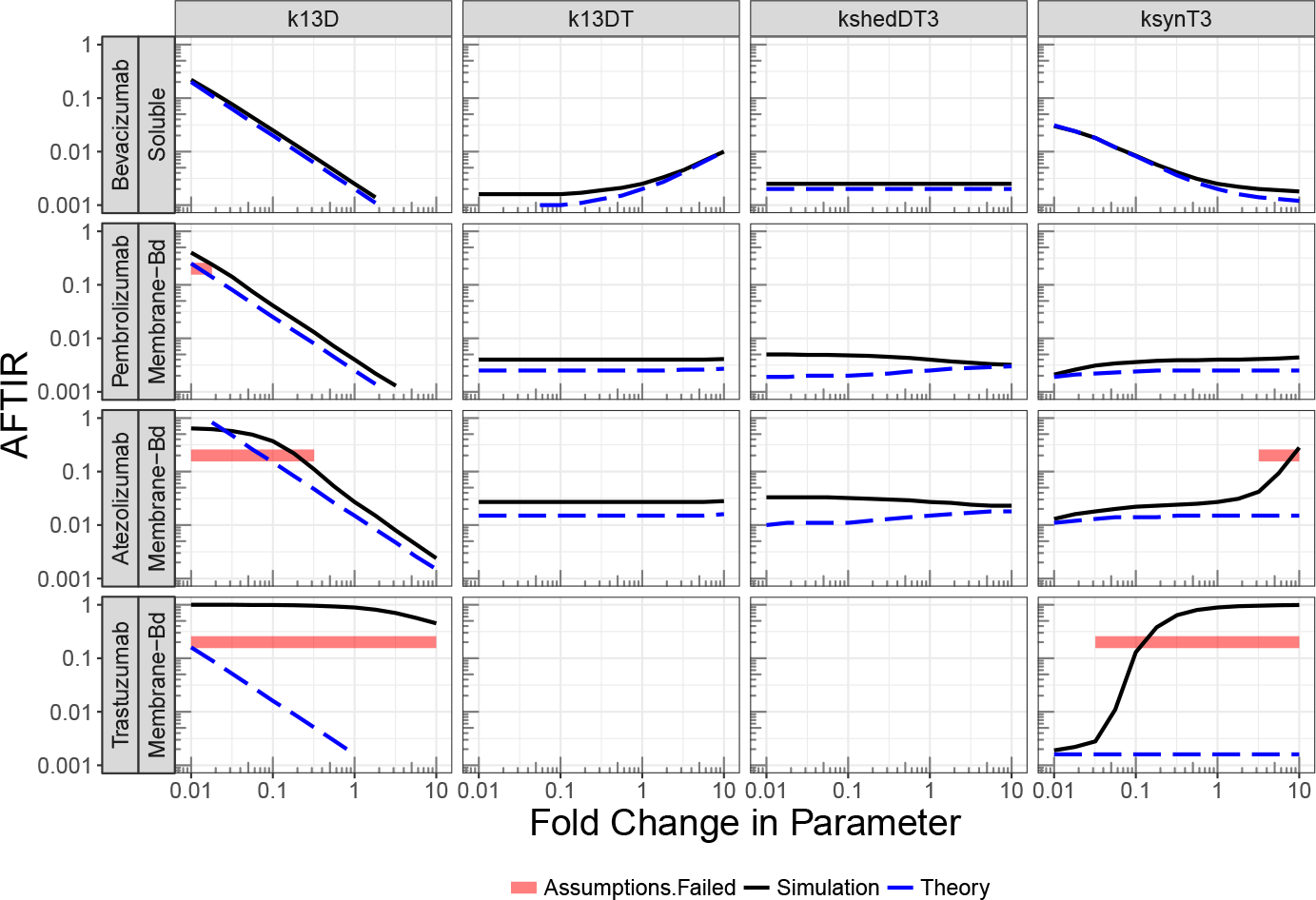
Parameter exploration where the parameter at the top of each column is changed from 0.01× to 10× from its baseline value. There is in general good agreement between theory and experiment, except for trastuzumab, when the target expression is high. Kss approximation.

**Fig. S3.**
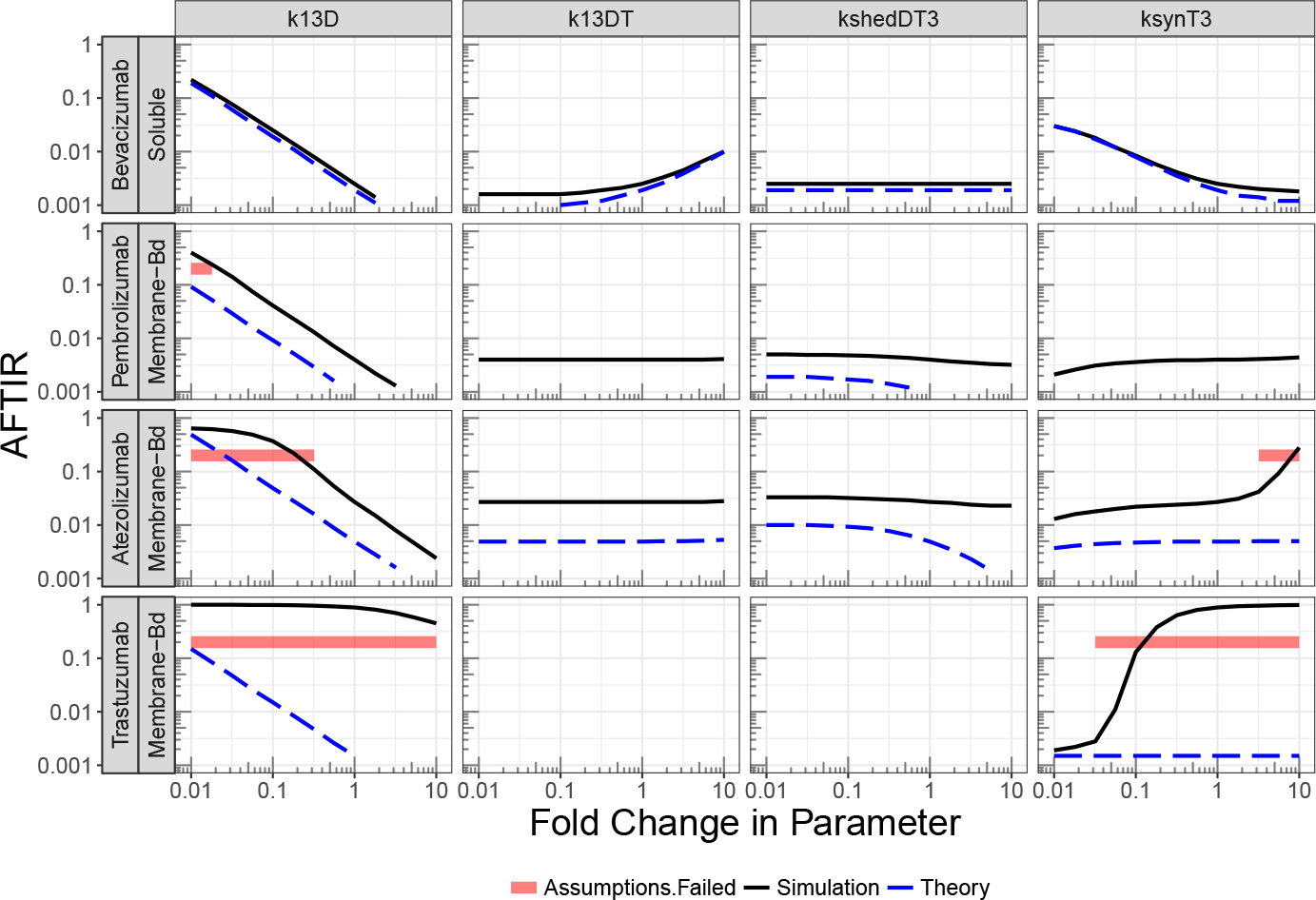
Parameter exploration where the parameter at the top of each column is changed from 0.01× to 10× from its baseline value. There is in general good agreement between theory and experiment, except for trastuzumab, when the target expression is high. Kd approximation.

### TFTIR simulations

**Fig. S4.**
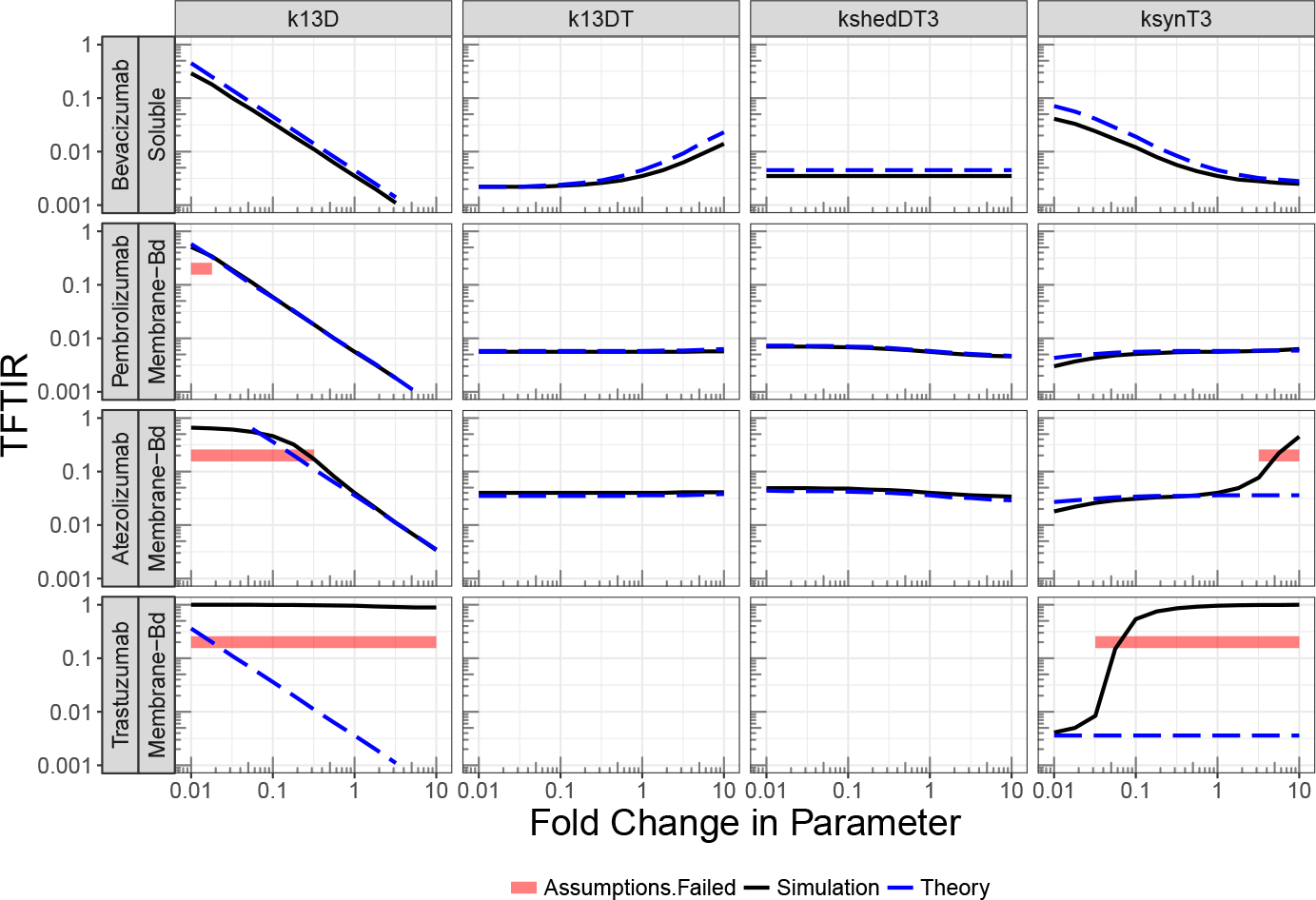
Parameter exploration where the parameter at the top of each column is changed from 0.01× to 10× from its baseline value. There is in general good agreement between theory and experiment, except for trastuzumab, when the target expression is high. Kssd approximation.

**Fig. S5.**
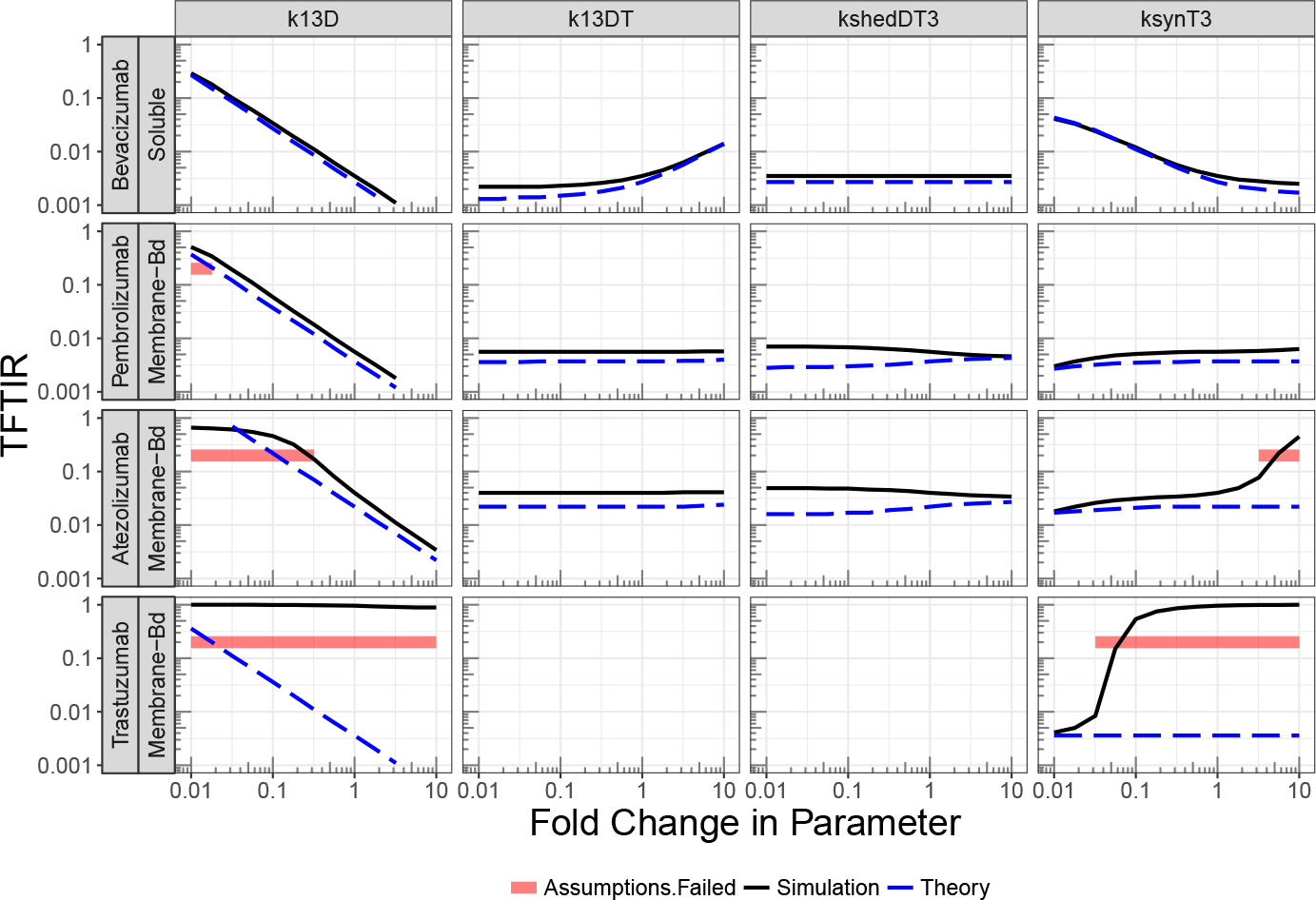
Parameter exploration where the parameter at the top of each column is changed from 0.01× to 10× from its baseline value. There is in general good agreement between theory and experiment, except for trastuzumab, when the target expression is high. Kss approximation.

**Fig. S6.**
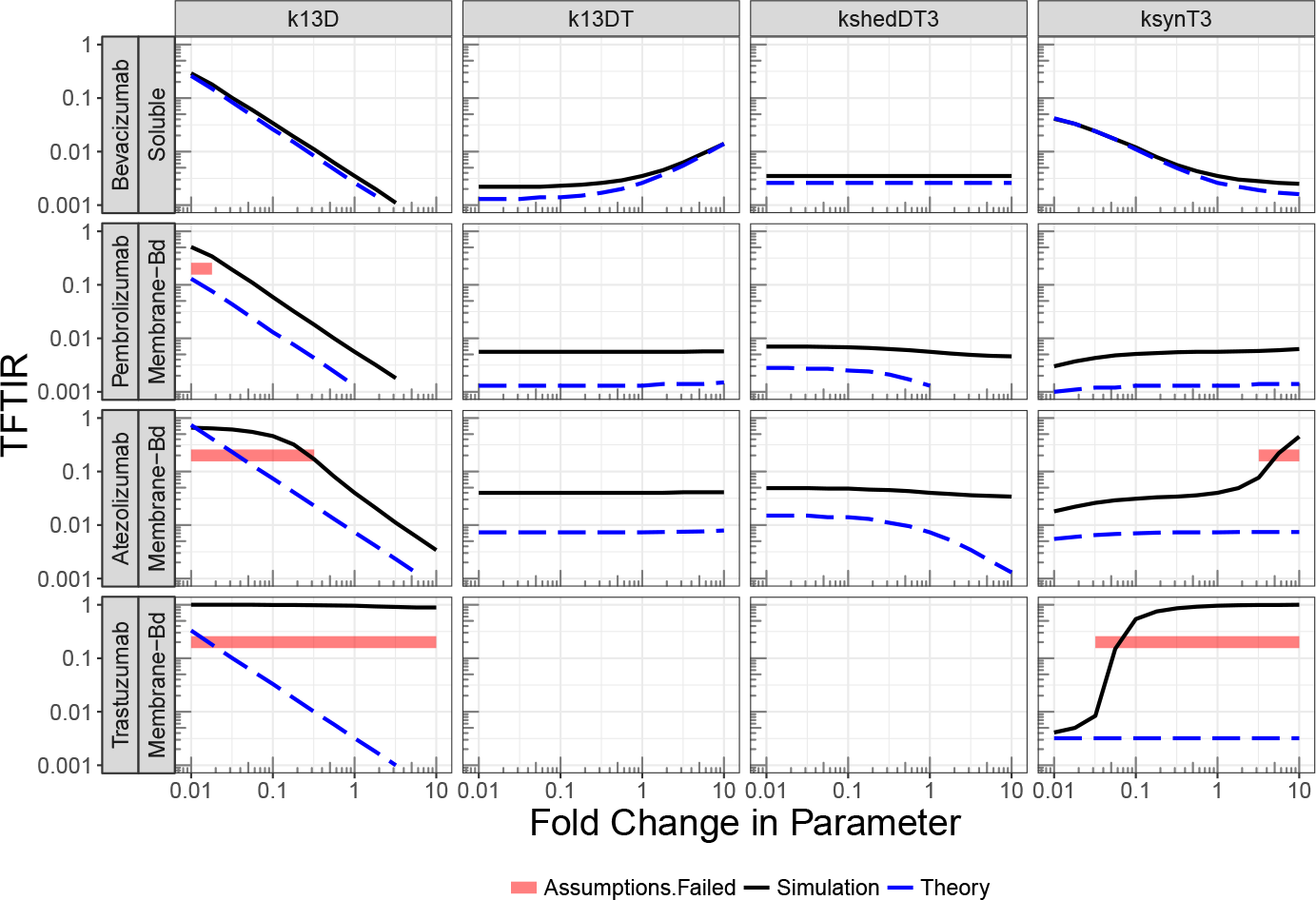
Parameter exploration where the parameter at the top of each column is changed from 0.01× to 10× from its baseline value. There is in general good agreement between theory and experiment, except for trastuzumab, when the target expression is high. Kd approximation.

